# Generative design approach to combine architected Voronoi foams with porous collagen scaffolds to create a tunable composite biomaterial

**DOI:** 10.1101/2023.09.05.556448

**Authors:** Marley J. Dewey, Raul Sun Han Chang, Andrey V. Nosatov, Katherine Janssen, Sarah J. Crotts, Scott J. Hollister, Brendan A.C. Harley

## Abstract

Regenerative biomaterials for musculoskeletal defects must address multi-scale mechanical challenges. We are developing biomaterials for craniomaxillofacial bone defects that are often large and irregularly shaped. These require close conformal contact between implant and defect margins to aid healing. While we have identified a mineralized collagen scaffold that promotes mesenchymal stem cell osteogenic differentiation *in vitro* and bone formation *in vivo,* its mechanical performance is insufficient for surgical translation. We report a generative design approach to create scaffold-mesh composites by embedding a macro-scale polymeric Voronoi mesh into the mineralized collagen scaffold. The mechanics of architected foam reinforced composites are defined by a rigorous predictive moduli equation. We show biphasic composites localize strain during loading. Further, planar and 3D mesh-scaffold composites can be rapidly shaped to aid conformal fitting. Voronoi-based composites overcome traditional porosity-mechanics relationship limits while enabling rapid shaping of regenerative implants to conformally fit complex defects unique for individual patients.

## 1. Introduction

Osseous defects of the skull occur secondary to trauma, congenital abnormalities, or after resection to treat stroke, cerebral aneurysms, or cancer [1-4]. Standard of care is calvarial reconstruction, while common (>35,000/yr in the US; including >10,000 cleft palate repairs)[5, 6], autologous bone reconstruction is a primary option [7], but has significant limitations including insufficient access to bone and donor site morbidity [8] as well as surgical complications (10-40% cases)[9-12]. There is significant intraoperative time required to shape autologous bone grafts to fit irregular defects, particularly for high risk defects (radiation, previous infection)[7]. While alloplastic materials (e.g., PEEK) are more easily shaped, they are plagued by complications such as extrusion and high infection rates (5-12 fold greater than autologous)[1, 13, 14]. A biomaterial implant that can be shaped precisely and quickly like an alloplastic implant but works like autologous bone would be transformative for craniofacial reconstruction.

We have reported a mineralized collagen scaffold for craniofacial bone repair. This porous scaffold endogenously activates Bone Morphogenic Protein (BMP) receptor signaling and promotes the expansion and osteogenic differentiation of rat, human, rabbit, and porcine mesenchymal stem cells (MSCs) without exogenous BMP [15-18]. It improves bone regeneration in rabbit craniofacial bone defects without the need for traditional osteogenic supplements [16]. However, while cell bioactivity and oxygen/nutrient biotransport in this biomaterial scale with material porosity, mechanical strength scales inversely with its porosity [7, 19]. The mechanical performance of the scaffold alone is insufficient for surgical handling and shaping required for large-scale clinical adoption. We developed scaffold-mesh composites by embedding a polymer mesh with millimeter-scale porosity into the collagen scaffold during fabrication [20-23]. These collagen-mesh composites display the osteogenic activity of the scaffold, the strength of the mesh, and can improve regenerative healing in a porcine mandibular defect [24]. However, poor conformal contact with the surrounding wound margin can inhibit healing [9, 25]. Common interventions such as resorbable plates do not reduce micromotion or improve conformal contact [26]. Given the irregular nature of craniofacial bone defects, approaches that facilitate rapid, intraoperative shaping to achieve close implant-defect conformal fitting are essential. We recently showed modifying mesh architecture can increase conformal contact in a cylindrical defect [23], but selective modifications based on prior knowledge of defect shape are not generalizable *de novo*.

Cellular (porous) materials are found broadly both in nature and in engineering applications where mechanical efficiency is essential [27, 28]. Generative design approaches inspired by these materials offer a route to develop three-dimensional [29] architectures with tailored structural and mechanical activity [30-32]. Cellular solids can be described using polyhedral unit cells and characterized by their relative density (ρ*/ρ_s_), the density of the porous structure (ρ*) normalized by the density of the solid wall (ρ_s_) [33, 34]. Open-celled pore structures are defined by the 3D organization of fibers termed struts. Low-density, open-cell foams display well-characterized mechanical performance metrics defined not by individual pore geometry but rather by the square of its relative density, (ρ*/ρ_s_) [33]. A well-defined linear elastic regime of these materials suggest an oversized implant may be reproducibly compressed then spring back to conformally fit a complex defect.

The decoupling of mechanical performance from pore architecture in low-density open-cell foams suggests a strategy to create foam-based meshes that retain their mechanical performance characteristics even after mesh trimming due to intraoperative shaping. Herein we propose a novel approach using architected meshes to create a scalable reinforcement strategy that allows post-fabrication shaping. Architected Voronoi foams are a subset of random, open-porous foams [35], defined by the random distribution of seed points in three-dimensional space which subsequently define the center of each pore of the Voronoi design (**Figure 1A**). This provides the opportunity to use generative design to define the density of seed point in 3D space rather than focus on the design of an explicit foam architecture. Their mechanical properties, like other low-density open-cell foams, are based on relative density, suggesting they can be controlled by the density of pores (which can be computationally defined) and the thickness of the struts that define the pores (which can be controlled by 3D printing; (**Figure 1B**). Recently *Martinez et al.* showed Voronoi structures 3D-printed with local variations in pore geometry and density to yield stiffer and more flexible regions [36]; Voronoi foams have been used as standalone materials to mimic the porous nature of bone [37]. However, their potential to form an adaptable mesh reinforcement in a composite biomaterial that can be shaped after fabrication has not been previously studied.

**Fig. 1.**
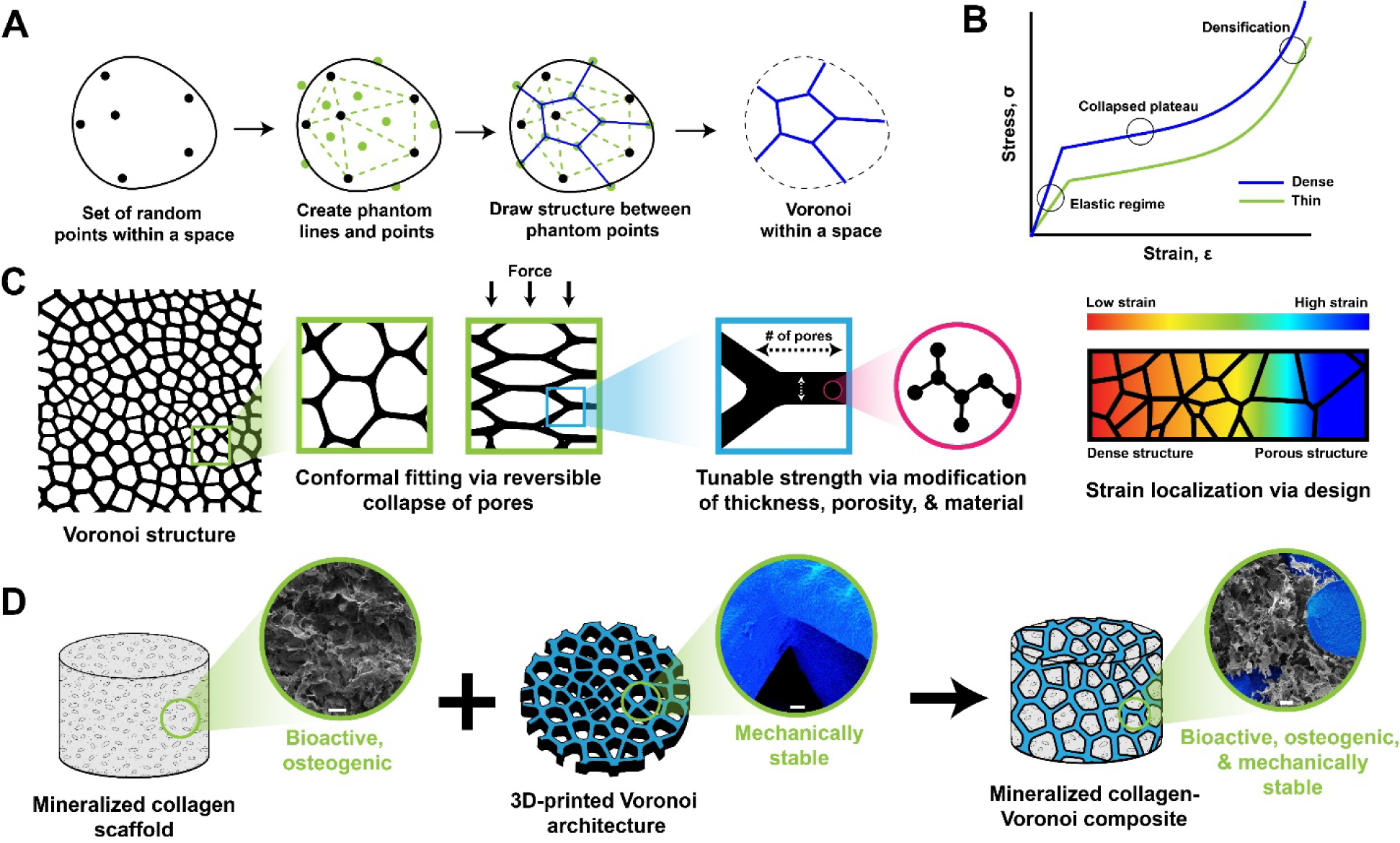
Tough and tunable Voronoi structures. (A) To create a Voronoi structure within any space, firstly, a set of “seed” points are added to the space. Then phantom lines (green) are drawn between these points, with phantom points (green) in the middle of these areas. Finally, the lines of the Voronoi structure (blue) are drawn between the phantom lines connecting the phantom points. (B) Representative stress-strain curve of foams, such as Voronoi structures. These structures are characterized by a linear elastic regime, a collapsed plateau, and densification. Increasing the density of Voronoi structures increases the modulus (elastic regime). (C) Voronoi structures can be applied to any size or shape and can achieve conformal fitting by the reversible collapse of pores. Additionally, the thickness of the individual struts in the architecture and the material properties can lead to tunable strength and elasticity. Finally, strain can easily be localized by printing designs with regions of variable porosity or density of material. (D) The goal of this study was to incorporate tunable Voronoi 3D-printed architectures into mineralized collagen scaffolds for use in bone repair. Mineralized collagen scaffolds offer excellent bioactivity and osteogenic properties due to the mineral, glycosaminoglycans, collagen, and porous nature of this material; however, this porosity is a detriment to mechanical robustness. Combining Voronoi architectures and mineralized collagen scaffolds offers a biomaterial to promote osteogenesis while also maintaining strength. Scale bar represents 100 µm.

In this article, we describe fabrication of a unique class of Voronoi-based polymeric reinforcement meshes, their integration into a mineralized collagen scaffold, and the functional performance of the resultant multi-scale composite (**Figure 1D)**. We show composite mechanical performance can be tailored via alterations to the Voronoi foam pore density and the strut thickness; moreover, we describe potential to leverage mechanical isotropy and predictive models for designing a Voronoi mesh to achieve a desired modulus (**Figure 1C)**. We describe biphasic Voronoi architectures to localize deformation in discrete regions of a composite. We show scaffold-mesh composites can be cut and trimmed to shape after fabrication and that inclusion of a Voronoi-mesh increases the conformal fitting capacity as measured via a push-out test. We also show Voronoi meshes can be fabricated from multiple polymers, demonstrating the broader bioengineering applicability of this concept. Distinct from designing a unique mesh geometry for each patient defect, we describe a strategy to create adaptable composites that can be shaped after fabrication to fit unique defect geometries without a priori knowledge of their shape. We hypothesize multi-scale composites will increase regeneration by limiting graft micromotion and improving cellular and vascular integration.

## 2. Methods

### 2.1 Fabrication of 3D-printed Voronoi structures

All three-dimensional Voronoi structures were designed using nTop software (nTopology, New York, USA) and two-dimensional (planar) designs were designed using Fusion360 (Autodesk, California, USA). Point spacing is defined as the distance (mm) between the randomly seeded points within a Voronoi structure.

#### Fabrication of photopolymerized Voronoi structures

Voronoi structures made of photopolymerized resin for porosity testing measured 10 mm cubes with three different porosities and were created by altering the Voronoi point spacing (4 mm point spacing, 3.5 mm point spacing, 3 mm point spacing) with 0.7 mm fiber diameter. The porosity was further minimized by printing 0.5 mm fiber diameter prints with 4 mm point spacing. These .stl files were printed using a Form 2 photopolymerization printer (Formlabs, Massachusetts, USA). A white standard resin was used to create all samples. Biphasic and uniform structures were made of white resin, and these designs were fabricated as 20 x 6 x 6 mm (length, width, height) Voronoi rectangular prints with a point spacing of 2 on one half of the structure and a point spacing of 3.75 on the other half, and printed with a 0.7 mm fiber diameter. These were compared to uniform Voronoi rectangles, measuring the same dimensions and a point spacing of 2 throughout the space.

#### Fabrication of laser-sintered Voronoi structures

Voronoi structures used for thickness testing and composite sheet fabrication were printed with laser sintered poly(caprolactone).[56-60] The Voronoi structures were manufactured using laser-sintering with a Formiga P110 (EOS, Krailling, Germany). These structures were fabricated using a blend of polycaprolactone, a bioresorbable and biodegradable polyester, with a 4% weight by weight ratio of hydroxyapatite, a naturally occurring mineral, used in previous studies with mineralized collagen scaffolds.[47, 48] Voronoi structures for thickness testing were fabricated as 25 mm cubes with the same pore spacing (3 mm) and various strut thicknesses (0.7 mm, 1 mm, 1.5 mm). Voronoi structures for composite sheet testing were fabricated as either 2D or 3D sheets, with 2D sheets measuring 74 x 74 mm squares with 245 cells and 80% cell scale, and 3D sheets measuring 74 x 74 x 6 mm (length, width, height) and 3 point spacing and 0.7mm strut thickness.

### 2.2 Fabrication of Voronoi-mineralized collagen composite structures

To fabricate mineralized collagen scaffolds and composites, a mineralized collagen suspension was created as previously described[39, 44, 46] and composites were fabricated via lyophilization similarly described previously [47, 48, 61]. Briefly, type I bovine collagen (Collagen Matrix, New Jersey, USA) was blended together with a mineral solution of phosphoric acid and calcium hydroxide (Sigma Aldrich), chondroitin sulfate (Spectrum Chemicals, New Jersey, USA), and calcium nitrate tetrahydrate (Sigma Aldrich) using a rotor-stator in a jacketed cooling vessel until well-blended[44, 46, 47]. To create mineralized collagen scaffolds, 24 mL or 48 mL of collagen suspension (for 3.5 mm or 7.5 mm thickness sheets, respectively) was pipetted into a 75 x 75 mm aluminum pan. Rectangular scaffolds for DIC testing were cut with a razor to size and cylindrical scaffolds for shape-fitting testing were cut with a 12 mm biopsy punch. To create Voronoi-mineralized collagen composites, collagen suspension was added to a 75 x 75 mm aluminum pan and Voronoi 3D-prints were carefully added to pan. The suspensions were then lyophilized using a Genesis freeze-dryer (VirTis, New York, USA) by dropping the temperature at a rate of 1°C/min to a 2 hour hold at −10°C. Resulting mineralized collagen scaffolds and composites were then solid, integrated structures.

### 2.3 Predictive equation for Voronoi structures

A predictive modulus equation for open-cell foams was used to predict the Elastic Modulus of Voronoi structures[34].

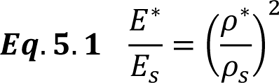

Where E* represents the Predictive Young’s Modulus, E_s_ represents the Young’s Modulus of the solid skeleton, and (p*/p_s_) represents the relative density. The predictive modulus values were determined by calculating the v/v% of Voronoi designs based on .stl files (known densities) and normalizing these to the design with the greatest volume of 3D-print (i.e. thickest or most dense print). Thus, the Voronoi architecture with the highest relative density was the “base” (value of 1) design, and was used to calculate the expected modulus of the remaining Voronoi architectures with smaller relative densities based on the known differences in relative density.

### 2.4 Compression testing of Voronoi structures

All Voronoi structures created underwent compression testing and were analyzed with a custom Matlab program in order to determine Young’s Modulus, Ultimate Stress, and Ultimate Strain.[34, 62] All structures were loaded onto platens facing the same direction (same face upward). To test isotropy, each design was tested on each face of the cube (x, y, z axis). Matching experimental data to a predictive equation was determined by normalizing the other designs tested to the design with the highest density of 3D-print.

Photopolymerized resin Voronoi structures were mechanically compressed by an Instron 5943 mechanical tester (Instron, Massachusetts, USA) with a 100 N load cell at a compression rate of 1 mm/min. Six samples were used for each design and each face tested. Laser-sintered polycaprolactone Voronoi structures printed as 25 mm cubes were mechanically loaded and unloaded to half the thickness of the print. An MTS Criterion model 43 (MTS, Minnesota, USA) was used to compress at a rate of 1 mm/sec with a 1 kN load cell and holding at 1 sec before releasing at a rate of 1 mm/sec. Videos of prints were taken using a Pixelink camera (Pixelink, Canada). Percent deformation of samples was determined by the difference in the strain after unloading compared to the strain at the start of the experiment. Eight samples were used for each thickness and face tested for 25 mm cube designs.

Biphasic Voronoi structures were compressed by an Instron 5943 mechanical tester (Instron) with a 100 N load cell at a rate of 3 mm/min to generate data for stress-strain curves (n=6). Scaffolds and composites were glued to 3D-printed bases (ABS) to stabilize these to the mechanical platens, with the dense region of the biphasic structure always glued to the base. Following embedding into the ABS base, scaffolds were speckle-patterned with waterproof India ink (BLICK Art Materials, Illinois, USA) using a gravity feed airbrush with a nozzle size of 0.3 mm (Got Hobby Inc., California, USA). During uniaxial compression testing, images were taken with a Canon EOS 5DS R DLSR camera and a Canon Macro 100-mm lens at a rate of 1 image every 5 seconds until an arbitrary point of maximum compression (Canon, Tokyo, Japan). The digital images were correlated using a previously described method of analysis[63]. Briefly, the sets of images taken during compression testing were correlated using a version of the MATLAB file package “Digital Image Correlation and Tracking” (Copyright © 2010, C. Eberl, D.S. Gianola, S. Bundschuh) modified by Elizabeth Jones (Improved Digital Image Correlation version 4 – Copyright © 2013, 2014, 2015 by Elizabeth Jones) to calculate local strain across scaffolds. For each set of scaffold images, the region of interest was set as the entire exposed scaffold. Reduced images were correlated first to generate initial guesses for displacements in full images and reduced image correlations were iterated 5 times (image reduction factor: 3, subset size: 221; threshold: 0.5; search zone: 3, grid step size: 10). Subsequently, full images were correlated using the reduced image correlation data and individually optimized subset sizes (subset size: 421-621; threshold 0.5; search zone: 2; grid step size: 20). Finally, displacements were smoothed prior to calculating strains to reduce noise (Gaussian distribution of weights; kernel size: 11; number of smoothing passes: 3; maximum size of contiguous non-correlated points to smooth over: 15) and local strains were calculated using a cubic (16-node) algorithm. Contour plots and line scans of strain could then be visualized. Line scan data was exported onto Microsoft Excel and average strain plotted as a function of position for each sample. Due to high displacements, images near the maximum compression did not correlate. Therefore, strain profiles were analyzed from the start of compression to simply show the localization of strain throughout the scaffold in biphasic versus uniform Voronoi structures.

2D and 3D Voronoi-mineralized collagen sheets underwent push-out testing to determine shape-fitting ability compared to mineralized collagen scaffolds without printed supports. 12 mm biopsy punches were used to remove samples from sheets, and these were added to two defect sizes in a Teflon mold, either 11.5 mm or 10.8 mm in diameter. The Teflon mold was applied to an apparatus allowing for a metal pin to push the composites or scaffolds through the defects; the same apparatus was used in previous work to test for shape-fitting ability[61]. The maximum push out force (N) was measured and compared between groups, with eight samples per group. 2D Voronoi composites were compared to mineralized collagen scaffolds of the same thickness (approximately 3.5 mm thick) and 3D Voronoi composites were compared to mineralized collagen scaffolds of the same thickness (approximately 7.5 mm thick).

### 2.5 SEM imaging of Voronoi and mineralized collagen composites

The 3D-print integration within mineralized collagen scaffolds was visualized via SEM of photopolymerized biphasic Voronoi-mineralized collagen composites, and 2D and 3D Voronoi-mineralized collagen composites. Biphasic composites were cut in both phases to expose the interior of the dense and porous regions, and 2D and 3D Voronoi composites were cut in half to visualize the center of these composites.

### 2.6 Statistics

Statistics followed testing outlined in literature and previous experiments [38, 64]. Data was first checked for normality (Shapiro-Wilk) and equal variance (Levene’s Test) of residuals, using a Grubb’s test to check for and remove any outliers. If data was normal with equal variance, a One-way ANOVA was used with a Tukey post-hoc to determine significance (p < 0.05) between more than 2 groups. For only two group comparisons, a two sample T-test was used, and data was marked only significant if the power was above 0.8.

## 3. Results

### 3.1 The mechanical performance of architected Voronoi foams can be defined by generative design algorithms but is dependent on print parameters

The mechanical performance of Voronoi foams created for scaffold reinforcement can be predicted based on the relative density of the mesh, which itself can be manipulated either by adjusting Voronoi seed point spacing or by changing the thickness of the printed struts. We created two homologous series of foams, adjusting the Voronoi foam architecture via the nTop software package (*design*) and strut thickness via the 3D printing process (*manufacture*).

#### Design

We first adjusted the Voronoi point spacing (3 – 4mm point spacing; 0.7mm fiber diameter), achieving a range of reproducible mesh relative densities (ρ*/ρ_s_): 19v/v%, 15v/v%, 12v/v% (**Figure 2A**). We also created a minimum density control, printing 0.5mm fiber diameter struts with a 4mm point spacing (6%v/v), though these structures were not mechanically stable. Voronoi structures (10mm cubes) were printed from .stl files using a Form 2 photopolymerization printer (Formlabs, Massachusetts, USA) using white standard resin, and were subsequently evaluated mechanically under unconfined compression. All groups displayed stress-strain relationships characteristic of open-cell foam structures, with defined linear elastic and collapse plateau regimes. As expected, the elastic modulus increased significantly with mesh density (**Figure 2A**, **Table 1, Supp. Table 1**). We subsequently evaluated the predictability of mesh Young’s Modulus as a function of relative density, calculating predicted changes in mesh moduli via known differences in mesh relative density. Experimental results of all three Voronoi structures (19v/v%, 15v/v%, 12v/v%) closely matched the predicted modulus for each group (**Figure 2A**). These results confirm generative design approaches to increase the relative spacing of Voronoi seed points reduces mesh density and results in a predictable decrease in mechanical performance of the printed mesh.

**Fig. 2.**
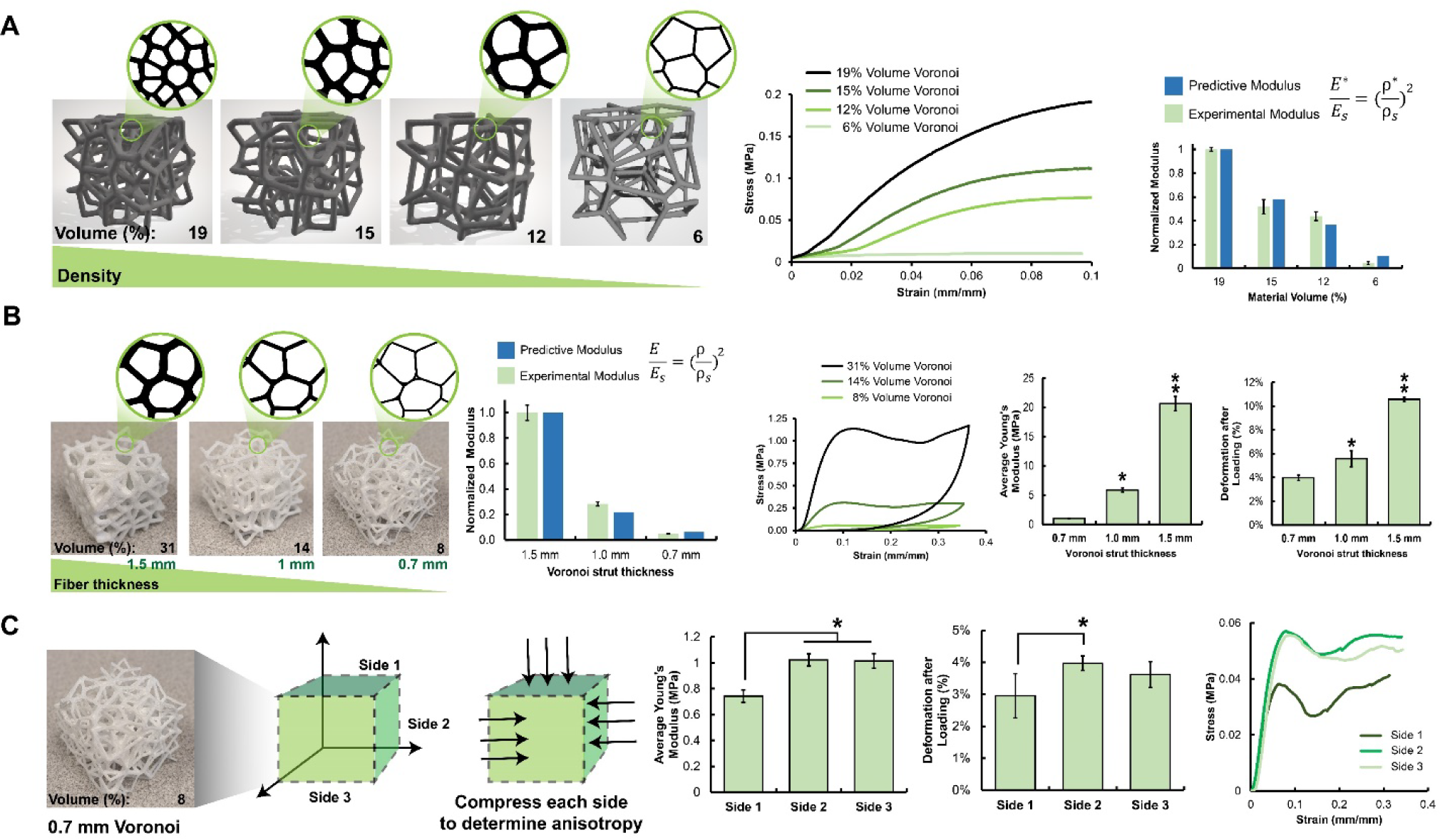
Tuning stiffness through Voronoi density and material thickness. (A) Four Voronoi designs (10 mm cubes) of different porosities altered by changing the pore architecture were fabricated with photopolymerized resin. Designs were compressed and the stress-strain profiles were recorded. The modulus of the four designs was compared to a predictive modulus for Voronoi architectures. (B) Three Voronoi architectures (25 mm cubes) with the same design and different strut thicknesses were printed out of polycaprolactone. The modulus of these designs was tested against a predictive modulus for Voronoi architectures. Samples were compressively loaded and unloaded and the stress-strain profiles were recorded. The average modulus and deformation after loading increased with increasing strut thickness. ** indicates the 1.5 mm group was significantly (p < 0.05) greater than all other groups. * indicates the 1.0 mm group was significantly (p < 0.05) greater than all other groups. (C) Isotropy was analyzed in the 0.7 mm thickness Voronoi designs printed with polycaprolactone and 8% volume material (25 mm cubes). Each of the sides of one Voronoi design (x, y, z axis) were compressively loaded and unloaded and the modulus and deformation after loading were examined. * indicates which group(s) were significantly (p < 0.05) greater than another group. Differences in the modulus between sides of the same design indicates anisotropy. Data expressed as average ± standard deviation (n=6).

**Table 1.** Determining predictive modulus and experimental modulus of Voronoi 3D-prints made of white resin based on point spacing and strut thickness.

| Predictive Equation |  |  |  |  | Experimental Data |  |
| --- | --- | --- | --- | --- | --- | --- |
| Design | v/v% | $\rho^*/\rho_s$ | $(\rho^*/\rho_s)^2$ | Normalized $E^*$ | Young's Modulus (MPa) | Normalized $E^*$ |
| 4 | 6.22 | 0.06 | 0.004 | 0.11 | $0.17 \pm 0.054$ | $0.05 \pm 0.01$ |
| 3 | 11.82 | 0.12 | 0.014 | 0.37 | $1.42 \pm 0.37$ | $0.40 \pm 0.04$ |
| 2 | 14.74 | 0.15 | 0.022 | 0.58 | $1.93 \pm 0.46$ | $0.54 \pm 0.06$ |
| 1 | 19.43 | 0.19 | 0.038 | 1.00 | $3.57 \pm 0.63$ | $1.00 \pm 0.02$ |

**Table 2.** Determination of predictive modulus and experimental modulus of Voronoi 3D prints made of polycaprolactone based on strut thickness.

| Predictive Equation |  |  |  |  | Experimental Data |  |
| --- | --- | --- | --- | --- | --- | --- |
| Design (mm) | v/v% | $\rho^*/\rho_s$ | $(\rho^*/\rho_s)^2$ | Normalized $E^*$ | Young's Modulus (MPa) | Normalized $E^*$ |
| 0.7 | 7.72 | 0.08 | 0.01 | 0.06 | $1.02 \pm 0.047$ | $0.05 \pm 0.002$ |
| 1 | 14.47 | 0.14 | 0.02 | 0.22 | $5.86 \pm 0.36$ | $0.28 \pm 0.017$ |
| 1.5 | 30.96 | 0.31 | 0.10 | 1.00 | 20.64 ± 1.23 | 1.00 ± 0.06 |

#### Manufacture

We subsequently adjusted mesh relative density using a consistent mesh design, but instead altering the thickness (0.5mm, 1.0mm, 1.5mm) of the printed struts. Large (25mm) cubes of Voronoi meshes were fabricated via a laser-sintered polycaprolactone process (**Figure 2B**), where increasing the strut thickness (0.7mm, 1mm, 1.5mm) increased mesh relative density (8%v/v, 14%v/v, 31%v/v). We again observed predictable increases in Voronoi foam mechanics, with well-defined linear elastic and collapse plateau regimes as well as elastic modulus scaling as (ρ*/ρ_s_)^2^ (**Figure 2B**, **Supp. Table 2, Supp. Fig. 1**). Elastic hysteresis, the ability for an over-sized implant to be compressed then spring back to fill space, plays an important role in conformal fitting capacity in complex defects [23, 38]. 3D-printed polycaprolactone meshes display significant deformation capacity with strut buckling and pore collapse phenotypes characteristic of low-density open-cell foams (**Supp. Video 1**). Predictably, while increasing strut thickness increased elastic modulus, this also led to greater permanent deformation and less elastic hysteresis. Videos of compressive loading and unloading of polycaprolactone 3D-prints demonstrated this change in flexibility, with the thinnest fiber print (0.7mm) showing buckling behavior useful for forming to fit a defect space without breaking.

### 3.2 Voronoi meshes display partial mechanical isotropy

Mechanical isotropy is an important consideration in the design of reinforcing meshes that display predictable properties [39]. We evaluated the isotropy of printed Voronoi (25mm laser-sintered polycaprolactone cubes; 0.7mm strut thickness; 8%v/v) foams. Each face of the Voronoi structure was compressed to evaluate axis-specific differences in stiffness and deformation. Voronoi foams demonstrated some degree of anisotropy with statistically significant differences in Young’s Modulus and deformation between some compression axes (**Fig. 2C**). Additional 10mm foam cubes fabricated using photopolymerizable resin (Formlabs) also displayed anisotropy (**Supp. Table 3**). While the random pore orientation of Voronoi foams should theoretically result in isotropic structures, isotropy requires numerous pores in each axis to remove effects of local morphology. So, while all printed designs demonstrated some degree of anisotropy at the scale at which they were tested (10mm and 25mm dia. cubes), moduli differences were on the order of a fraction of 1MPa and deformation differences of 1%, which may not be significant during application of Voronoi structures clinically.

### 3.3 Local changes to Voronoi mesh architecture enable localized control over composite deformation

While an isotropic Voronoi foam would support uniform deformation and mechanical properties, it may equally be important to concentrate deformation to distinct regions (e.g., near the edge of an implant to facilitate conformal contact). We fabricated biphasic Voronoi meshes comprised of porous and dense regions (Porous: 3.75 point spacing, 13v/v%, 0.7mm print; Dense: 2 point spacing, 36v/v%, 0.7mm print) with a continuous interface. We created collagen-mesh composites formed from *Biphasic (porous and dense regions)* or *Uniform (porous region only)* Voronoi meshes fabricated from white photopolymerized resin (Formlabs). Following lyophilization, Voronoi meshes were fully embedded into mineralized collagen scaffolds. We observed some voids around mesh elements near the center of dense Voronoi meshes (**Fig. 3A**), suggesting an upper limit (ρ*/ρs:0.36) to the density of Voronoi meshes that could be embedded into scaffolds. Unlike continuous displacement profiles exhibited for uniform Voronoi meshes (**Supp. Video 1**), Biphasic Voronoi meshes displayed a stress-strain profile that suggested the presence of unique *linear elastic*–*collapse plateau*–*densification* regions for first the Porous mesh compartment followed by the Dense mesh compartment after the full collapse of the Porous compartment (**Fig. 3B**). A similar biphasic stress-strain profile was visible in the Voronoi-mineralized collagen composites, with incorporation of the mineralized collagen increasing the Young’s Modulus of both the porous and dense regions compared to the 3D-mesh alone (**Supp. Table 4, Supp. Fig. 2A,B,D**). The predictive capacity of the local moduli of a biphasic Voronoi design was much less accurate than for monophasic Voronoi meshes (**Supp. Fig. 2C**). Digital Image Correlation (DIC) analyses confirmed that while uniform strain distributions were observed across Uniform scaffold-mesh composites, strain was regionally concentrated in composites formed from a Biphasic mesh (**Fig. 3C**). Strain was localized to the Porous zone of the Biphasic mesh and at a higher level than in composites formed from the Uniform mesh (**Fig. 3D**).

**Fig. 3.**
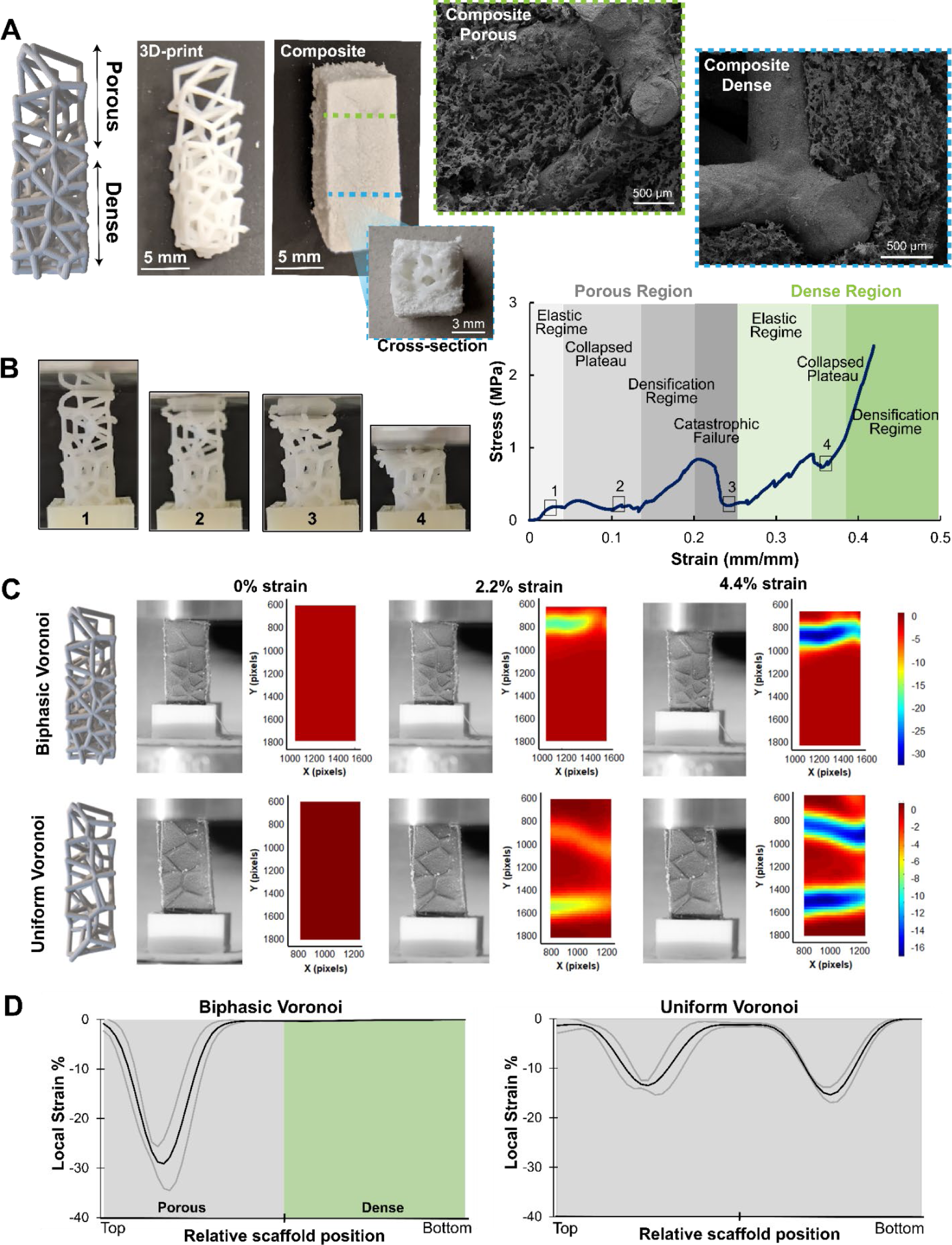
Localizing strain through biphasic Voronoi design. (A) Biphasic Voronoi designs were 3D-printed with photopolymerized resin into designs with a distinctly porous region and dense region. These were combined with mineralized collagen scaffolds to create a composite material. SEM images demonstrate the integration of the 3D-print with surrounding mineralized collagen in the dense region (blue) and porous region (green). (B) Biphasic Voronoi 3D-prints were compressed and the stress-strain profile was recorded. The stress-strain profile was comprised of two different curves, whereas the porous region first compressed until failure (grey), then the dense region was compressed (green), outlined on the stress-strain profile with matching images of the 3D-print under compression. (C) Biphasic Voronoi and mineralized collagen composites were compared to Uniform Voronoi mineralized collagen composites. These were compared under compression and Digital Image Correlation to map strain regions and create contour plots. At 0%, 2.2%, and 4.4% global applied strain, images of the composite are alongside representative strain localization images, with blue representing the greatest strain. (D) Line scans of the strain on biphasic Voronoi composites and uniform Voronoi composites were created from Digital Image Correlation data. Average strain at a global applied 4.4% strain was plotted as a function of position for each sample, with the average (black line) and standard deviation (interval contained by gray lines) of strain magnitude along the entire length of each scaffold. The porous region is demonstrated by a grey background and the dense region with green (n=6).

### 3.4 Planar scaffold-mesh composites fabricated using two-dimensional Voronoi meshes can be readily shaped post-fabrication

Planar composites may be particularly valuable for some cranial bone and orbital repair applications. We fabricated planar composite sheets by embedding a planar polycaprolactone Voronoi mesh into mineralized collagen scaffolds (**Fig. 4A, Supp. Fig. 3**). Voronoi planar sheets were printed of polycaprolactone and measured 74×74mm with 245 cells and 80% cell scale (Fusion360, Autodesk software). Sheets of resultant scaffold-mesh composite could be readily shaped post-fabrication with scissors or a razor blade (**Supp. Video 2**). We biopsied 12mm diameter cylindrical specimens from a large composite sheet, then tested the push-out force from model cylindrical defects (11.5mm or 10.8mm). Results were compared to mineralized collagen scaffolds alone. There was no difference in push-out force between the planar Voronoi composite versus the mineralized collagen scaffold. Further, there was no increased force required to push the mineralized collagen scaffold or the planar Voronoi composite through the 10.8mm defect versus the larger 11.5mm defect, suggesting out of plane bending of the planar reinforcement (**Figure 4B**). While planar scaffold-mesh composites formed from a planar Voronoi mesh were able to be trimmed to fit and increased mechanical strength, this design did not provide additional conformal fitting capacity.

**Fig. 4.**
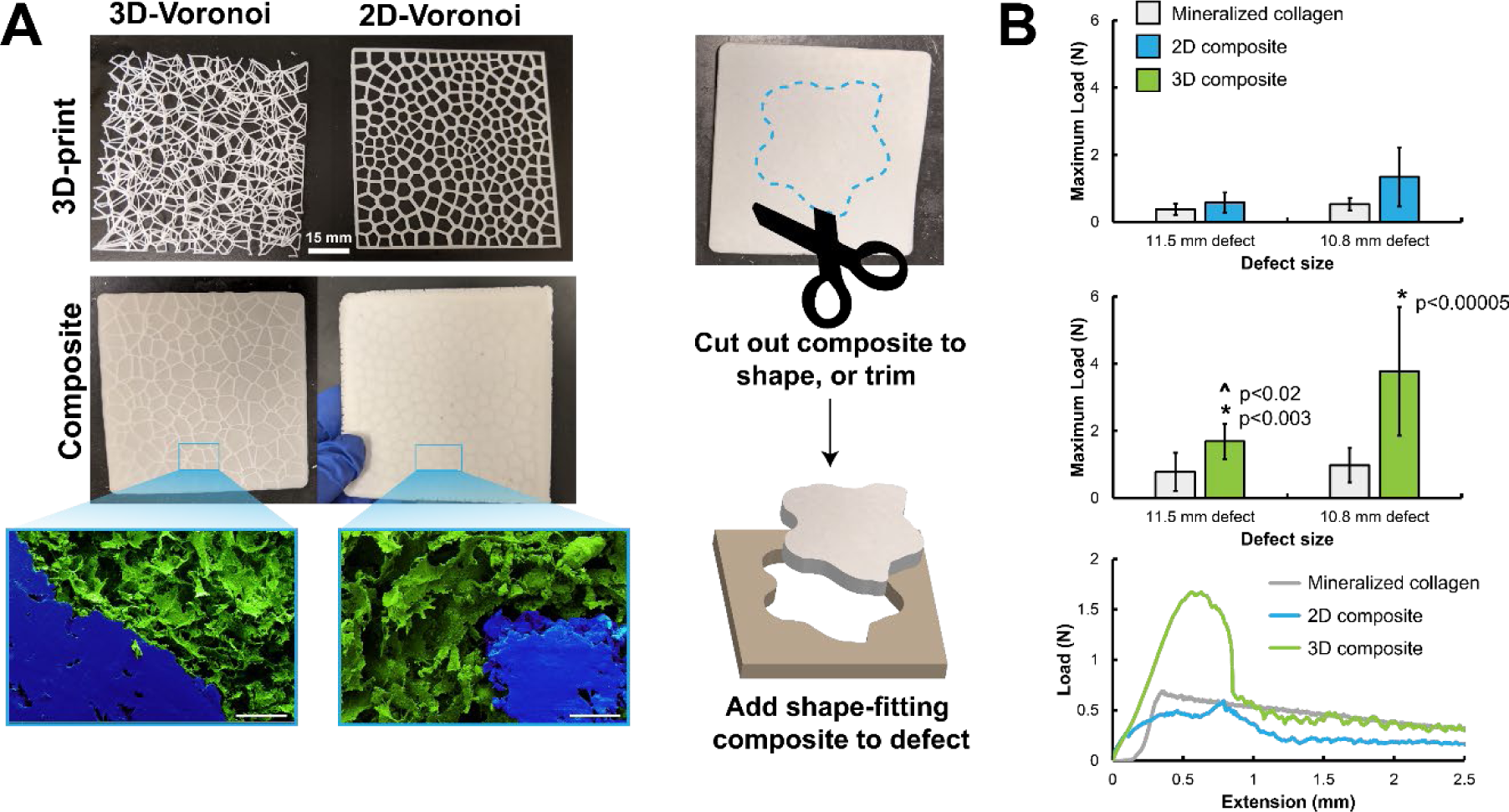
2D and 3D Voronoi composite sheets for shape-fitting applications. (A) Polycaprolactone was 3D-printed in a 75 x 75 x 3 mm sheet as a 3D-Voronoi design and printed as a 75 x 75 x 1 mm sheets as a 2D-Voronoi design. Designs were lyophilized with mineralized collagen to create 3D and 2D Voronoi composites. False-colored SEM images of the internal structure of the composites demonstrate good integration of the 3D-print (blue) with surrounding mineralized collagen (green). Scale bar represents 250µm. Composite sheets were fabricated for handlers to be able to cut a desired shape with scissors or a razor blade or trim an existing shape and fit this into the defect space via the conformal fitting capacity of Voronoi structures. (B) Shape-fitting analysis of 12mm biopsies of 2D and 3D Voronoi-mineralized collagen composite sheets (2D composite, 3D composite, respectively) compared to mineralized collagen scaffolds. Biopsies were added to a push-out testing device with two different sized defect spaces (11.5 mm and 10.8 mm) and the maximum load required to force design through the defect spaces was recorded and demonstrated in the Load vs. Extension graph. If a material has shape-fitting properties it will require a greater load in the smaller defect space (10.8 mm). * indicates significant difference between the 3D Voronoi composite and the mineralized collagen scaffold at the respective defect size. ^ indicates significant difference between the 3D Voronoi composite between the two defect sizes. Data expressed as average ± standard deviation (n=8).

### 3.5 Large scaffold-mesh composites fabricated using three-dimensional Voronoi meshes can be readily shaped post-fabrication and increase conformal fitting

Large three-dimensional composites may be valuable for aspects of facial bone repair applications (e.g., mandibular repair). We fabricated large (74mm x 74mm x 6mm, 3 point spacing, 0.7mm strut thickness) Voronoi composites by combining 3D polycaprolactone Voronoi meshes with mineralized collagen scaffolds to create a composite thicker than planar Voronoi structures (**Figure 4A**). Like planar composites, these fully three-dimensional Voronoi composites could be trimmed and shaped with scissors or a razor blade post-fabrication while maintaining structural integrity (**Supp. Video 2**). We again assess conformal-fitting ability for 12mm diameter (by 6mm thickness) specimens cut from a larger scaffold-mesh composite using a biopsy punch in a manner identical as that for planar composites. Unlike the planar Voronoi composites, the 3D-Voronoi composites required significantly greater force to push through 11.5mm defects and 10.8mm defects than did the mineralized collagen scaffold control group. Additionally, pushing the 3D-Voronoi composite through the smaller 10.8mm defect required significantly greater force than the larger 11.5mm defect, indicating 3D-Voronoi designs biopsied from a sheet could still retain shape-fitting ability even after deformation of the whole.

## 4. Discussion

The successful repair of CMF defects involves biomaterial implants that balance osteogenesis and mechanical integrity. Complications can arise from mismatched host bone and biomaterial properties, such as stress-shielding and micromotion. Stress-shielding can increase surrounding bone resorption, while micromotion (28-150µm of device movement) can lead to fibrous tissue formation around the implant and reduced bone formation [40]. While mineralized collagen scaffolds has a history of improved bone regeneration [41-46]; challenges with macroscale mechanical performance and fitting complex defects [47, 48] motivate this work to develop a scalable scaffold composite. We created a mineralized collagen 3D-print composite reinforced with a tunable Voronoi architecture to precisely control biomaterial mechanics to avoid implant rejection due to mechanical properties.

Voronoi mesh reinforcement reported here offers several advantages over more prescribed reinforcement approaches recently described. Notably, Voronoi structures offer a unique density-moduli relationship that decouples mechanical performance from explicit foam geometry. We hypothesized that matching experimental porosities to predictive moduli could demonstrate the unique tailorable nature of these architectures. Our results show it is possible to create a homologous series of Voronoi meshes, independent of mesh composition, and achieve predictable changes in mesh mechanical performance. Further, our results demonstrate mechanical performance can be jointly defined by mesh architecture and print parameters (e.g., print thickness), with smaller thickness prints (0.7mm) able to undergo reversible deformation, useful for surgical handling, defect fitting, and mechanical integrity.

Generally, the moduli of these Voronoi structures were far from the moduli of the softest portion of bone (cancellous 0.1-2GPa), as the densest Voronoi structure tested (∼31v/v% PCL) achieved a modulus of 20MPa [49]. Although all structures tested had moduli less than that of cancellous bone, designs printed with 0.7mm fiber thickness all had greater moduli than dry mineralized collagen scaffolds. Exactly achieving the moduli of host bone may not be necessary for effective bone repair, as soft mineralized collagen scaffolds have demonstrated the ability to regenerate rabbit calvarial defects with a much lower modulus when hydrated (kPa)[50, 51]. Further, small changes in stiffness, on the order of hundreds of kilopascals, has demonstrated changes in bone regeneration; the addition of crosslinking mineralized collagen scaffolds has led to increases in bone formation [52]. We believe even a small increase in stiffness may be able to improve bone formation in these collagen scaffolds, and limiting the 3D-print polymer volume to <30v/v% may avoid large releases of acidic byproducts which can cause a persistent inflammatory responses [53].

Scaffold anisotropy may add complexity to surgical handling. Due to the random structure of Voronoi designs, these by nature should be isotropic; however, in all our tests we noticed some degree of anisotropy. This could be attributed to the low ratio of size to point spacing (i.e. 10mm cubes with 4mm point spacing). We hypothesize that a larger design, such as sheets of Voronoi structures, would result in a more isotropic structure, and CMF defects are typically 25mm or larger in size. Voronoi computation models relating moduli to predictive equations have used a minimum of 27 cells to accurately assess isotropy [54], suggesting the differences in modulus may be due to an insufficient number of pores in that axis of compression.

Fabricating a biomaterial with various regions of stiffness is desirable for conformal-fitting applications, allowing for radial compression and inner stability. To evaluate this, we designed a biphasic 3D-print with a porous and dense region and successfully combined this with mineralized collagen scaffolds to create a composite material, and further compared this to a composite with only one reinforced phase (porous). The compression of the biphasic 3D-print and composite demonstrated stress-strain profiles of two different density materials, and DIC analysis demonstrated that strain is solely concentrated at the porous region. We successfully created a biphasic composite that could increase the moduli of mineralized collagen scaffolds while also offering tunable strain profiling by incorporating regions of different 3D-print porosity architecture, allowing additional control over mechanical design.

Lastly, we investigated the potential for Voronoi reinforced composites to aid surgical practicality. While the end goal was to fabricate Voronoi mineralized collagen composite “sheets” that could be cut and shaped intraoperatively and subsequently press-fit into defects, here we explored the ability for Voronoi-reinforced composites to increase push-out force in a set of standard sized cylindrical defects as a proxy for conformal fitting. We fabricated planar (2D) and 3D variants of Voronoi structures to create composites, 2D-Voronoi targeted for thinner cranial defects (i.e. rabbit skull) and 3D-Voronoi targeted for thicker defects (i.e. porcine or human jaw). Due to the large porosity of these designs, mineralized collagen suspension was able to fill in and around the pores, creating uniform composite structures. These were easily cut to a desired shape and size post-fabrication with scissors or a razor blade. While they could be press-fit into different diameter defects (highlighting our composite approach aids rapid fitting of complex geometries), 2D-Voronoi composites did little to add to shape-fitting due to the radial stiffness of these designs. However, 3D-Voronoi composites were shapeable and resulted in higher push-out forces than mineralized collagen scaffolds and higher forces required to push through smaller defect spaces, indicating shape-fitting ability and the potential to minimize micromotion *in vivo*. The lack of shape-fitting ability in 2D composites could be due to the thickness of the Voronoi fibers and the lack of a three-dimensional compressible network, which allows for more buckling without irreversible pore collapse. As a result, ongoing efforts will be seeking to explore alternative design variants of more 2D-reinforcing geometries as there is a significant need for such a material [55].

Critically, this work establishes that a standard predictive equation [34] used to model the mechanical performance of a wide variety of porous materials could be used to describe 3D printed Voronoi structures. Even small density designs (<8v/v%) could add additional stiffness to mineralized collagen scaffolds. We demonstrated that these designs could be fabricated with two different densities to localize strain, and these could be printed as sheets for easier surgical handling. We emphasize that there are many considerations for applications of these Voronoi reinforcements, such as lower bounds of print thickness (0.5mm strut thickness), upper bounds to avoid acidic byproduct release during degradation (>30%), and print porosity which may limit microscale polymer infiltration into print spaces (>2 point spacing). Although thinner prints provide a smaller increase in stiffness of the overall 3D-print, we believe the addition of shape-fitting ability due to print flexibility would improve osteogenesis over an increase in stiffness. Future work will investigate osteogenesis and osseointegration of 2D and 3D-Voronoi sheet composites compared to mineralized collagen scaffolds. We plan to investigate the ability of these structures to form new bone in rabbit calvarial defects, and how shape-fitting ability influences the integration and push-out strength required after implantation and subsequent bone formation. Overall, our work presents a highly tailorable reinforcing support with control of material thickness, flexibility and stiffness, conformal fitting and region-specific mechanical properties, and translation to any size and shape defect.

## 5. Conclusion

A critical challenge to craniomaxillofacial bone repair is the design of implants with osteogenic and mechanical properties to promote successful new bone formation. Due to the nature and size of these defects, improper implant fit to the defect space can prevent new bone formation from occurring. We examined the addition of tailorable 3D-printed Voronoi structures, or open-porous architectures, to mineralized collagen scaffolds. We demonstrate that the moduli can be tailored and predicted based on print porosity and thickness, biphasic designs can be developed to localize strain profiles for enhanced mechanical control, and composite structures can be fabricated in large sheets for easy shaping and cutting to fit complex defects. Additionally, we demonstrate that 3D Voronoi structures within collagen scaffolds can conformally fit to defect spaces, allowing for compression of structures to better fit within non-uniform shaped defects. These composite materials represent a scaleable, novel biomaterial with enhanced tunability for overcoming the challenge of poor implant fit, which ultimately leads to repair failure, of craniomaxillofacial defects.

## Acknowledgements

We would like to thank the following facilities at the University of Illinois for the ability to complete this work: Carl R Woese Institute for Genomic Biology, the High Temperature Materials Laboratory, and Roger Adams Laboratory. We would also like to thank the following for assisting with completion of this work: Peter Kurath (UIUC) for assistance with compression testing, Angela Andrada (Harley Lab) for assisting with compressing testing, and Aidan Gilchrist (Harley Lab) for assistance with Matlab codes to analyze compression data, and the remainder of the Harley Lab for their support. This work was supported by the Office of the Assistant Secretary of Defense for Health Affairs Broad Agency Announcement for Extramural Medical Research through the Award No. W81XWH-16-1-0566. Research reported in this publication was also supported by the National Institute of Dental and Craniofacial Research of the National Institutes of Health under Award Number R21 DE026582 and R01 DE030491. We are also grateful for funds provided by the NSF Graduate Research Fellowship DGE-1144245 (MJD) to perform this research. The interpretations and conclusions presented are those of the authors and are not necessarily endorsed by the Department of Defense, NIH, or NSF.

## Contributions (CRediT: Contributor Roles Taxonomy[65, 66])

**Marley J. Dewey:** Conceptualization, Data curation, Formal Analysis, Visualization, Investigation, Methodology, Writing – original draft, Writing – review & editing. **Raul Sun Han Chang:** Investigation, Data curation, Formal Analysis, Visualization, Writing – review & editing. **Andrey V. Nosatov:** Investigation, Formal Analysis, Writing – review & editing. **Katherine Janssen:** Investigation, Methodology. **Sarah J. Crotts:** Investigation, Methodology. **Scott J. Hollister:** Conceptualization, Resources, Supervision, Writing – review & editing. **Brendan Harley:** Conceptualization, Resources, Project administration, Funding acquisition, Supervision, Writing – review & editing.

## Data Availability

All raw and processed data will be accessible through Mendeley Data DOI: 10.17632/sdn9z5hc6g.1, which upon publication will be published.

## Notes

### Competing Interest Statement

The authors have declared no competing interest.

