## Supplementary Data for "Generative design approach to combine architected Voronoi foams with porous collagen scaffolds to create a tunable composite biomaterial"

^1^ Dept. of Materials Science and Engineering

^2^ Dept. Chemical and Biomolecular Engineering

^3^ Carl R. Woese Institute for Genomic Biology

^5^ Cancer Center at Illinois

University of Illinois at Urbana-Champaign

Urbana, IL 61801

^4^ Dept. of Biomedical Engineering

Georgia Institute of Technology

Atlanta, GA 30332

**Corresponding Author:**

B.A.C. Harley

Dept. of Chemical and Biomolecular Engineering

Carl R. Woese Institute for Genomic Biology

Cancer Center at Illinois

University of Illinois at Urbana-Champaign

110 Roger Adams Laboratory

600 S. Mathews Ave.

Urbana, IL 61801

**Supplementary Table 1.** Evaluating the Young’s modulus of Voronoi structures printed with photopolymerizable resin (n=6). Voronoi structures were printed with using either 0.5 or 0.7 mm diameter thicknesses, with varying pore spacing (4, 3.5, 3 mm) to create a library of porous materials. The print volume for the entire print was based on a shape volume of 10 mm^3^.

| **Sample** | **Side** | Young's Modulus (MPa) | Print volume (v/v%) |
| --- | --- | --- | --- |
| 0.5mm dia. 4 pore | 1 | 0.13 ± 0.05 | 6.22 |
|  | 2 | 0.18 ± 0.04 |  |
|  | 3 | 0.09 ± 0.07 |  |
| 0.7mm dia. 4 pore | 1 | 1.98 ± 0.16 | 11.82 |
|  | 2 | 1.35 ± 0.17 |  |
|  | 3 | 1.00 ± 0.18 |  |
| 0.7mm dia. 3.5 pore | 1 | 2.35 ± 0.26 | 14.74 |
|  | 2 | 2.39 ± 0.16 |  |
|  | 3 | 1.65 ± 0.23 |  |
| 0.7mm dia. 3 pore | 1 | 4.35 ± 0.07 | 19.43 |
|  | 2 | 3.49 ± 0.44 |  |
|  | 3 | 2.51 ± 0.33 |  |

**Supplementary Table 2.** Young’s Modulus, Ultimate Stress and Strain, as well as deformation after loading were evaluated for polycaprolactone Voronoi structures with different fiber thicknesses (0.7, 1, and 1.5 mm). Additionally, the 0.7 mm cube design was compressed on each of its 3 faces (x, y, z) to evaluate anisotropy (side 1, 2, 3).

| **Sample** | **Young's Modulus (MPa)** | **Ultimate Stress (MPa)** | **Ultimate Strain (mm/mm)** | **Deformation after Loading (%)** |
| --- | --- | --- | --- | --- |
| 0.7 mm | 1.02 ± 0.05 | 0.05 ± 0.003 | 0.09 ± 0.004 | 3.97 ± 0.23 |
| 1.0 mm | 5.86 ± 0.36 | 0.31 ± 0.01 | 0.10 ± 0.002 | 5.58 ± 0.68 |
| 1.5 mm | 20.64 ± 1.23 | 1.15 ± 0.07 | 0.12 ± 0.002 | 10.57 ± 0.18 |
| Side 1 | 0.74 ± 0.05 | 0.04 ± 0.003 | 0.07 ± 0.006 | 2.95 ± 0.69 |
| Side 2 | 1.02 ± 0.05 | 0.05 ± 0.003 | 0.09 ± 0.004 | 3.97 ± 0.23 |
| Side 3 | 1.01 ± 0.06 | 0.05 ± 0.004 | 0.09 ± 0.002 | 3.61 ± 0.41 |

**Supplementary Table 3.** Voronoi structures printed with photopolymerized White resin were compressed along three orthogonal axes to evaluate anisotropy. *: p < 0.05 different Young’s Moduli between compression axes.

| **Design** | **Side** | 1 | 2 | 3 |
| --- | --- | --- | --- | --- |
| 0.5 mm dia. 4 pore | 1 |  |  |  |
|  | 2 |  |  |  |
|  | 3 |  |  |  |
| 0.7mm dia. 4 pore | 1 |  | * | * |
|  | 2 | * |  | * |
|  | 3 | * | * |  |
| 0.7mm dia. 3.5 pore | 1 |  |  | * |
|  | 2 |  |  | * |
|  | 3 | * | * |  |
| 0.7mm dia. 3 pore | 1 |  | * | * |
|  | 2 | * |  |  |
|  | 3 | * |  |  |

**Supplementary Table 4.** The Young’s Modulus, Ultimate Stress, and Ultimate Strain of mineralized collagen scaffolds, biphasic Voronoi-scaffold composites, and biphasic Voronoi 3D-prints alone. While biphasic Voronoi 3D-prints (and the resultant composites) contain a dense and porous region to locally alter mechanical response, values were determined based on the shape of the stress-strain curve regions for open-cell foam structures (see Figure 3B and Supplementary Figure 2B,D).

| **Group** | **Young's Modulus (MPa)** | **Ultimate Stress (MPa)** | **Ultimate Strain (mm/mm)** |
| --- | --- | --- | --- |
| Scaffold | 0.31 ± 0.14 | N/A | N/A |
| 3D-Print (Porous) | 7.36 ± 2.48 | 0.24 ± 0.06 | 0.07 ± 0.03 |
| 3D-Print (Dense) | 13.69 ± 2.65 |  |  |
| Composite (Porous) | 10.82 ± 1.53 | 0.17 ± 0.01 | 0.04 ± 0.01 |
| Composite (Dense) | 31.33 ± 3.76 |  |  |

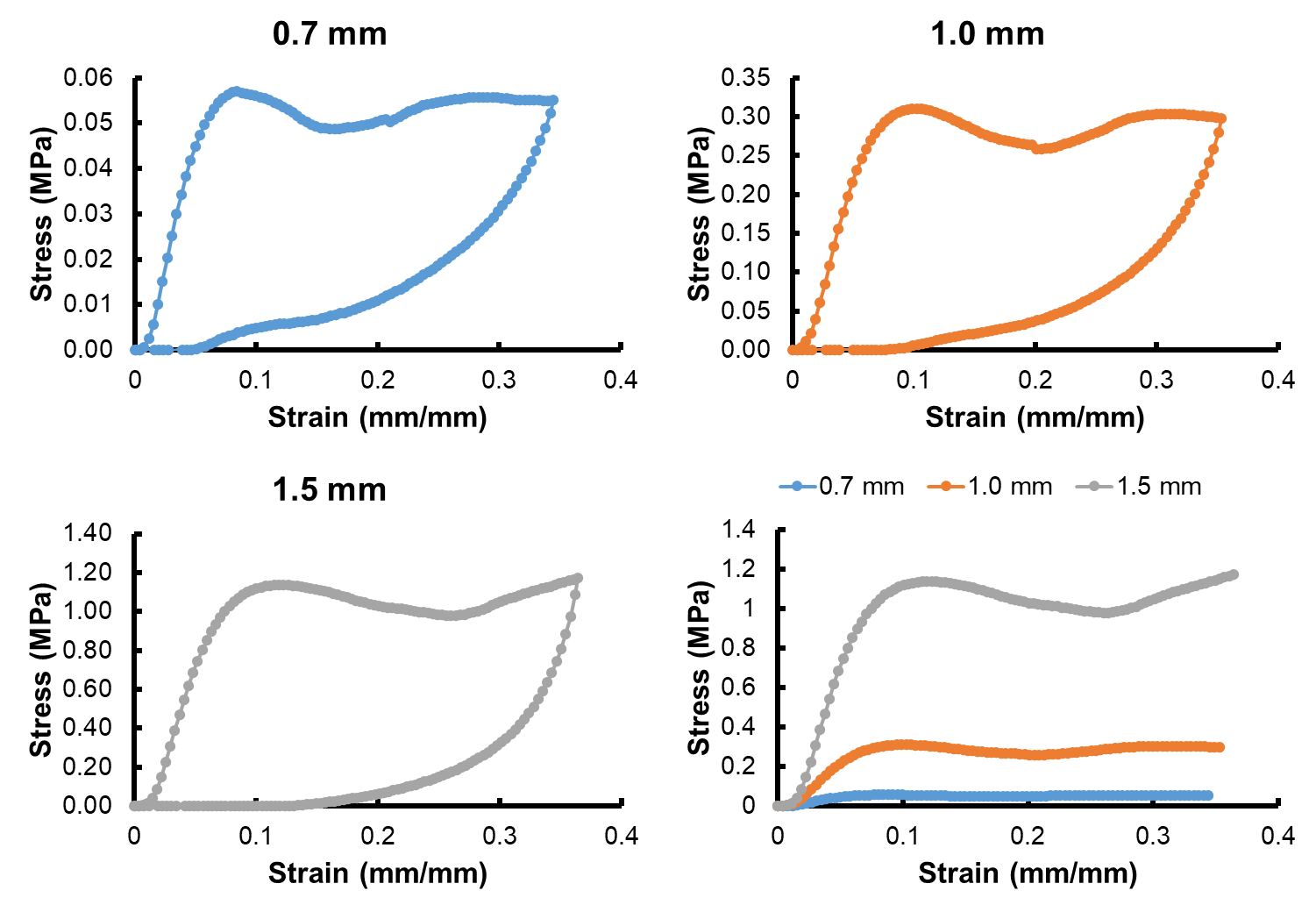

**Supplementary Figure 1.** Voronoi structures made of polycaprolactone were printed with different fiber thicknesses (0.7, 1, and 1.5 mm) and representative compressive loading and unloading curves were produced. The right-most graph is a representative stress-strain curve of laser-sintered PCL Voronoi cubes of different thicknesses overlaid on the same graph under loading only.

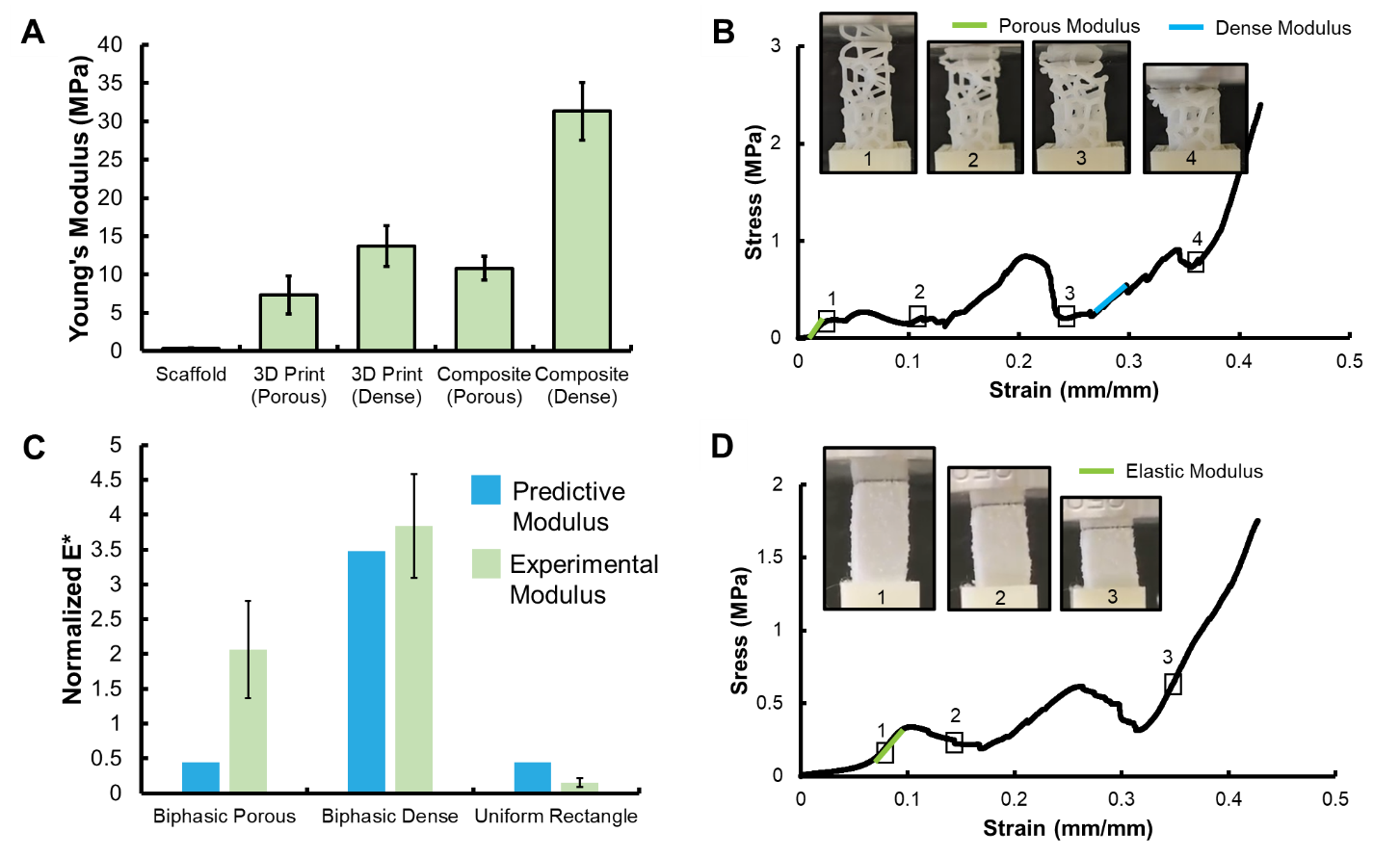

**Supplementary Figure 2.** Mechanics of mineralized collagen scaffolds reinforced with a biphasic Voronoi 3D-print (Objective 2). (A) Young’s Modulus of mineralized collagen scaffold, the porous and dense region of the biphasic Voronoi 3D-print, the porous and dense region of the mineralized collagen scaffold reinforced with a biphasic Voronoi 3D-print. All groups are significantly different (p < 0.05) from each other. Data expressed as average ± standard deviation (n=8). (B) Representative stress-strain curve of biphasic Voronoi 3D-prints. (C) Predictive and experimental modulus (E*) of rectangular Voronoi 3D-Prints. The normalized Young’s modulus of the porous and dense region of the biphasic print and the Young’s modulus of a uniform rectangular Voronoi print were all compared to the predictive normalized moduli based on cube Voronoi structures printed with the same material (see Figure 3B and Supplementary Table 4 (n=8)). (D) Representative stress-strain curve of mineralized collagen reinforced with biphasic Voronoi 3D-print.

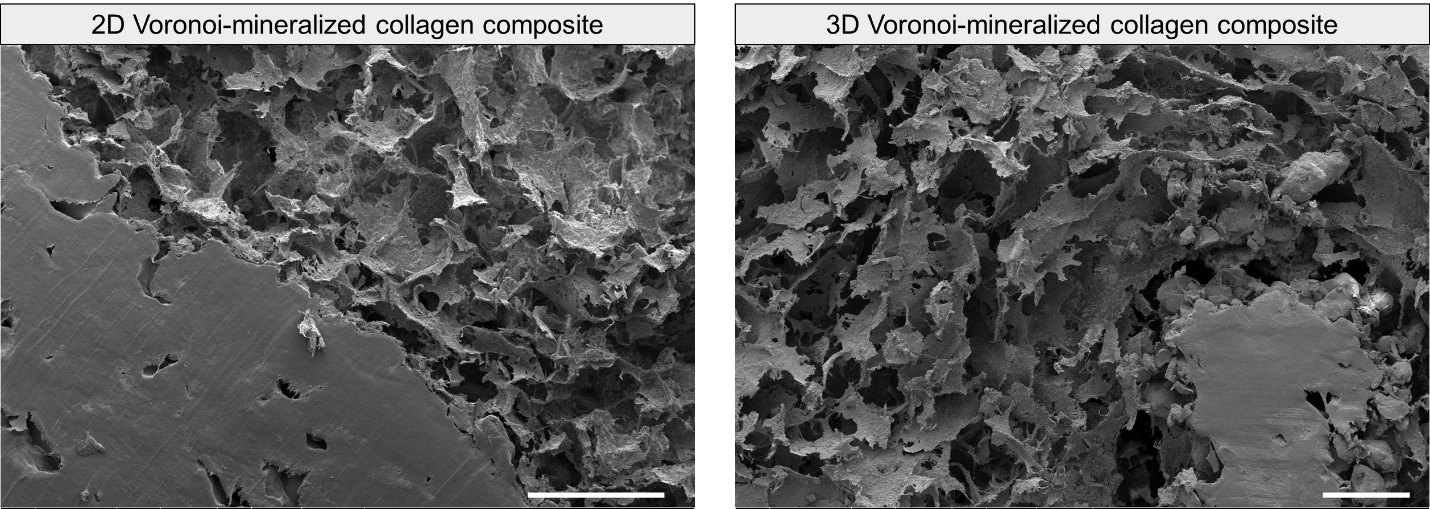

**Supplementary Figure 3.** SEM images demonstrating integration of PCL 3D-print and mineralized collagen in 2D and 3D Voronoi-mineralized collagen composites. Scale bar represents 250 µm. False-colored image used in Figure 4A.

**Supplementary Video 1.** Videos showing laser-sintered PCL Voronoi structures with increasing strut thicknesses (0.7, 1.0, 1.5 mm) under compressive loading and unloading. Fiber breaks are visible in the thicker diameter cubes (1.5 mm). Videos can be viewed at the following location in the folder title “Large PCL cubes of different thicknesses- loading and unloading_Supp. Video 1”: <https://uofi.box.com/s/tkx3l3stnjb87m1yzuh71chuxhs4idv2>.

**Supplementary Video 2.** Video depicting cutting and shaping 2D and 3D Voronoi mineralized collagen composite sheets with scissors or a razor blade. Videos can be viewed at the following location in folder titled “Cutting 2D and 3D Voronoi composite sheets_Supp. Video 2”:

<https://uofi.box.com/s/tkx3l3stnjb87m1yzuh71chuxhs4idv2>.
